# FRET-based aptamer assay for sensitive detection of *Salmonella paratyphi* A and revealing its molecular interaction with DNA gyrase

**DOI:** 10.1101/2020.04.02.021881

**Authors:** RM Renuka, Nikhil Maroli, J Achuth, K Kadirvelu

## Abstract

Rapid pathogen detection and identification of its serovars are crucial to provide essential treatment during pandemic circumstances. Herein, we developed a facile and versatile FRET-based aptasensor for rapid *Salmonella paratyphi* A detection. The ssDNA aptamers specific towards pathogenic *Salmonella paratyphi* A were generated via whole-cell SELEX. The aptamer was conjugated onto quantum dot (QD) that served as the molecular beacon and graphene oxide (GO) was used as fluorescence quencher. The detection of *Salmonella paratyphi* A leads to the quenching of QD fluorescence due to the non-covalent interaction between GO and CdTe quantum dot. The assay shows a detection limit up to 10 cfu·mL−1 with no cross-reactivity towards closely related species. The spiking analysis demonstrated an inter-assay coefficient of variance less than 8 % and recovery rate between 85%-102% mitigates assay reliability. Further analysis with commercially available ELISA kit validated the reliability of the developed aptasensor. Furthermore, molecular dynamics simulation was used to establish the mechanism of action of generated aptamer against bacterial DNA gyrase protein. The strong non-bonded interaction energies along with hydrogen bonds between the aptamer and protein inhibit the function of the bacteria.

## Introduction

*Salmonella* is one of the common bacterial pathogens responsible for typhoid fever, septicaemia, gastroenteritis, and other associated diseases. The untreated infection of this pathogen leads to death and severe organ damages such as liver and kidney failure. Salmonella bacterial pathogens are one of the unattended infections that led to death, in many geographic areas, particularly in developing countries^1–2^. Consumption of contaminated poultry, eggs, raw meat, dairy products, and chronic asymptomatic carriers are the major source of the pathogens^3^. Salmonella enterica serovars Typhi has long been recognized as the key causative agent for enteric fever, whereas, Paratyphi (A, B, C) accounted for the insignificantproportion of incident rates^4–6^. Among the *S. enterica* serovar Paratyphi, the prevalence of *Paratyphi* A in southeast Asia is prominent and accounts for 30 to 50% of enteric fever cases [14]. Being common food-borne bacteria, *Salmonella paratyphi* A remains as main cause of typhoid fever along with conditions like enter gastritis, sicchasia and food poisoning in humans and animals. Several data showthe rising consequence of enteric fever caused by S. paratyphidue to theincreased drug resistance ^7–10^ The emergence of multidrug-resistant strains andinsignificant cross-protection of *Salmonella* serovar Typhi vaccine administration are the major reason for S. paratyphi prevalence^11–14^.

Conventional detection techniques of Salmonella strains rely on culture and serological tests, which is time-consuming (3-6 days) and labor-intensive due to their substantially low specificity and sensitivity. Also, inadequacy of employing enriched culture lies in its inability to distinguish between viable-but-non-culturable (VBNC) cells^15^.Molecular biology-based techniques are easy to implement and high-throughput potential application such as polymerase chain reaction and multiplex PCR^16^, real-time PCR^17–19^, and loop-mediated isothermal amplification (LAMP) ^20–21^ are widely used. However, the nucleic acid-based molecular detection assays are incapable of distinguishing between viable or dead cells, since DNA persists in the environment extensively beyond the loss of cell viability^22^. Also, feasibility of mRNA-based detection is technically demanding and degradation through RNase could produce false-negative results^23^. An immunoassay approach is more specific as well as proficient in establishing a qualitative and quantitative determination of antigen. Nevertheless, these are labile to falsepositive results due to cross-reactive antibodies (polyclonal) and protein aggregation^24^.

Aptamers are high-affinity synthetic nucleic acid molecules generated against any target of choice from a combinatorial DNA library through an *in vitro* selection process termed SELEX ^25, 26^. With the potential to substitute antibodies, aptamers have become an attractive option as molecular affinities. Apart from its considerate affinity and stability, aptamers hold advantages like ease of synthesis and modification, cost-effective free of degradation and denaturation. Unfortunately, only very few aptamers targeting *Salmonella paratyphi A* have been generated till date and among these, none has tremendously increased the assay sensitivity without employing pre-enrichment strategies. Also, various aptasensors based on quartz crystal microbalance ^27^, electrochemistry^28^, Surface Enhanced Raman Spectroscopy (SERS)^29^ and fluorescence^30^based platforms efficiency principall yhave been reported, fluorescence-based platforms remain the preferred method owing to their superior sensitivity and reproducibility. Graphene oxide (GO) recently emerged as the most intriguing nonmaterial for sensing applications. GO has exceptional competence towards biomolecule interaction and together with its compatibility in solution make them as the material of choice in bio-sensing platforms^31^. FRET-based platforms efficiency principally relies on the donor and acceptor pair’s selection. GO with its remarkable electronic properties can serve as excellent energy acceptor by quenching the fluorescence of dyes in its excited states^32^. Quantum dots having broad absorption and narrow emission spectra together with negligible photo-degradationthan conventional dyes, also QD’s are an excellent source of electron donors in FRET-related assays. QDs can retain high fluorescence intensities for hours in contrast to organic dyes through continuous excitation. The bacterial cells labeled with QDs yield more fluorescence when equated with organic dyes (fluorescein isothiocyanate) ^33^. QDs are extremely sensitive to even a small change in donor-acceptor distance and thereby serve as the key context of FRET.

In this study we have developed a fluorescence-based aptasensor platform for *Salmonella paratyphi A* detection. The aptamers generation was carried out through whole-cell SELEX and it is characterized for its selectivity and affinity through Aptamer Linked Immunosorbent Assay (ELASA). Aptamer with better limit of detection and stable structure (sal1)was designated as a candidate for platform development.Sal1 was conjugated with CdTeQDs and allowed to interact with GO successively. Fluorescence quenching is enabled upon binding of aptamer labelled QDs with GO through π-π interaction. A target invoked fluorescence restoration has been observed in the presence of *Salmonella paratyphi A*. Further, the developed aptasensor was validated onto various standard cultures and artificially spiked matrices for assessment for its efficacy.

## Materials and Methods

### 2.1 Chemicals

All the salts were procured from Hi-media Labs, India and bacteriological culture media from BD-Difco. The aptamer library, primers, and biotinylated probes were synthesized at 1 mM and 25 nM scales, respectively at IDT, USA. The stock and working dilutions of the library and the primers were prepared in MilliQ water. Taq DNA polymerase and PCR buffers were purchased from Sigma, India. Streptavidin-horseradish peroxidises (HRP) were procured from Sigma, USA. pGEM-T vector and T4 DNA ligase for cloning were purchased from Promega, USA. Streptavidin, Graphite Cadmium oxide, Tellurium, Thioglycolic acid (TGA), N-hydroxysuccinimide (NHS), 1-(3-Dimethylaminopropyl)-3-ethylcarbodiimidehydrochloride (EDC) were procured from Sigma Aldrich, USA.

### 2.2 Bacterial strains and culture conditions

All bacterial cultures were procured from American Type Culture Collection (ATCC), USA. *Salmonella paratyphi A* (ATCC 9150) was used as target strain and other bacterial cultures *Klebsiella pneumonia* (ATCC 12022), *Salmonella typhimurium* (ATCC 14028), *Pseudomonas aeroginosa* (ATCC 27853), *Shigella sonneii* (ATCC 25931) were utilized as control strains. Cultures were grown at 37 °C in LB broth, harvested at log phase and serially diluted to requisite concentrations (10 cells) with Phosphate buffered saline (PBS pH7.4).

### 2.3 DNA library and primers

Random ssDNA library (nmol) was synthesized and purified by polyacrylamide gel electrophoresis (PAGE). The DNA pool consists of 87-base central random region oligonucleotides flanked by 22nt and 25nt constant sequences at the 3’ and 5’ ends that acted as primer binding regions for amplification. The biotin-labelled reverse primer was used for the aptamers blotting assays. Sequence details of primers employed and aptamers generated from present study were included in **table 1**.

**Table 1:**
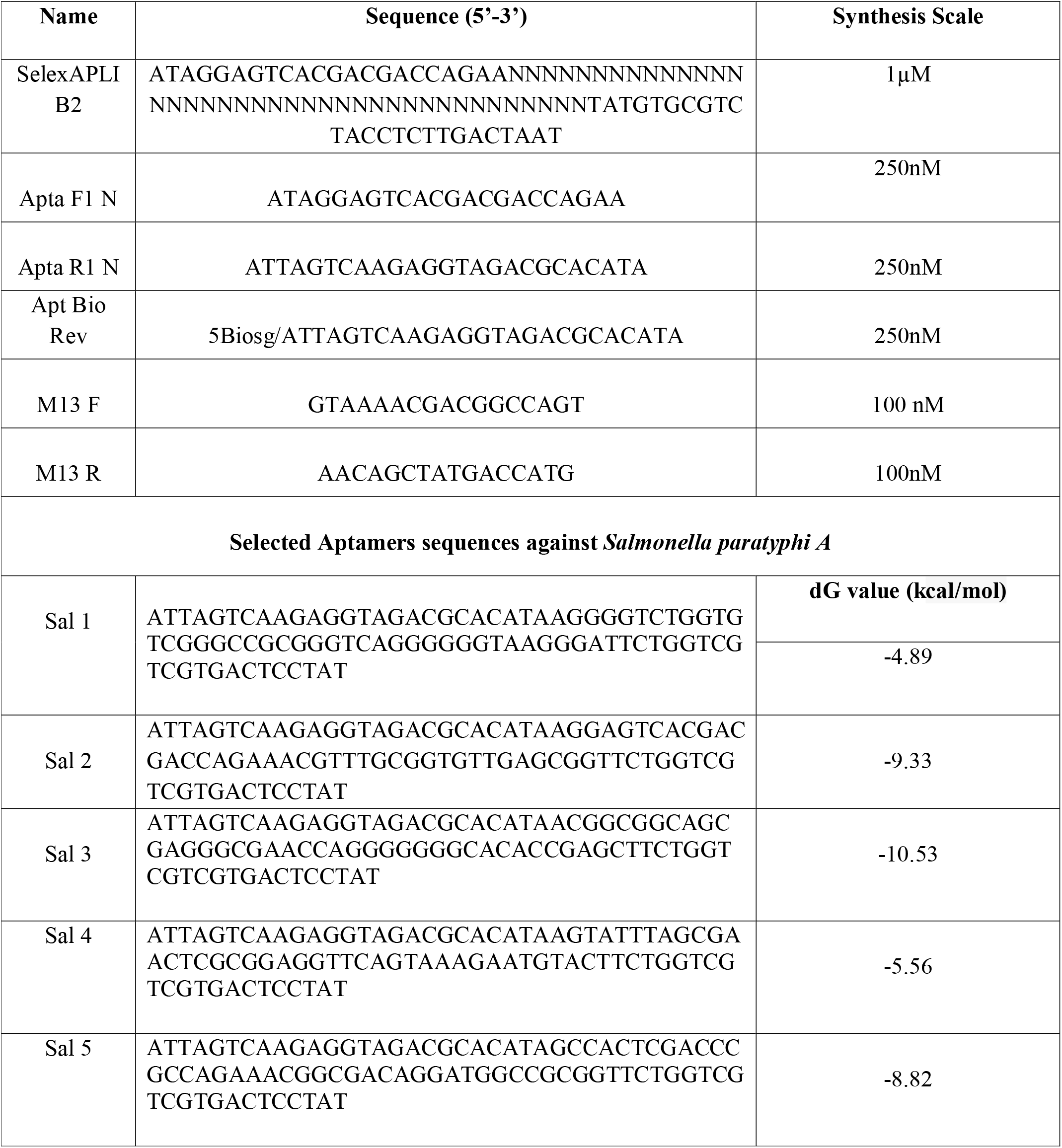
List of primers used for aptamers and sequence determined by sequencing.

### 2.4 Aptamer generation and characterization

#### 2.4.1. Whole-cell SELEX

*Salmonella paratyphi A* specific aptamers were generated through whole bacterial cell-SELEX as reported in our previous study ^34^ In short,200 nmol of the ssDNA pool and *Salmonella paratyphi A* (10^7^ cfu.mL^-1^) were incubated for 60 min at room temperature in binding buffer (100mM NaCl, 2mM KCl, 5mM MgCl_2_, 2mM CaCl_2_, and 20mM Tris HCl (pH 7.4)) supplemented with 0.1 % bovine serum albumin(BSA)to reduce nonspecific binding. Unbound aptamers fractions were removed through successive washing steps. The cell-boundaptamers were eluted with 1x PCR buffer through heat denaturation (95° C for 5 min) followed by snap freezing (on ice for 5 min) and centrifuged at 10,000 rpm for 5 mins. The collected supernatant fraction was enriched through PCR and product quality was assessed by agarose gel electrophoresis. A total of eleven rounds of selection were performed; seven rounds were directed towards *Salmonella paratyphi* A and four rounds of counter or negative selection towards *Salmonella typhimurium, Shigella flexnerii, E.coli* O157:H7 and *Yersinia entreocolitica* cells as negative selection counterparts.

#### 2.4.2. Cloning of selected aptamer sequences

Followed by selection cycles, the final reactive aptamer pool was subjected to poly A tailing and cloned into pGEMT vector (Promega USA). The transformed colonies were confirmed through colony PCR screening method with T7 F and T7 R primers. These clones were then subjected to plasmid extraction and subsequently preceded for sequence determination. Possible secondary structures and its corresponding Gibbs free energy values were predicted with the help of M-fold software^35^. Quadruplex forming G-rich sequences (QGRS) of generated aptamers were predicted using QGRS Mapper^36^.

#### 2.4.3. Characterization of *Salmonella paratyphi* A aptamers: Specificity and Sensitivity assay

The aptamers (Sal1 and Sal2) sensitivity against *Salmonella paratyphi A* was determined by Indirect ELASA (aptamer linked immuno sorbent assay). Briefly, 96 well microtiter plate (Genetix, India) was coated with 10^7^ cfu.mL^-1^ of *Salmonella paratyphi A* in 100 μL PBS (pH 7.4) at 4°C overnight. The unbound sites were blocked with 3% BSA in PBS (w/v) at 45 °C for 1h and washed thrice with both PBST and PBS. The biotinylated aptamers were subjected to heat denaturation at 95°C for 10 min and flash cooled on ice. Subsequently, various dilutions (1:500 to 1:64000) of aptamers in PBS was added into the microtiter plates and incubated for 1 h at 37°C. Followed by PBST and PBS washing, 1:2000 streptavidin-HRP conjugate was added and incubated for 1 hour at 37°C. With stringent washing, plates were developed with O-phenylenediamine (OPD; Sigma, US) and the reaction was stopped with 30 μL of 3N H_2_SO_4_. Absorbance values were recorded at 450nm using Biotek multi-mode plate reader. Initial library pool was used as control. Further, the limit of detection of generated aptamers was determined through indirect ELASA. Different *Salmonella paratyphi* A concentrations (10 to 10 CFU mL) were coated onto microtiter plates and were proceeded with ELASA as mentioned above. To determine the specificity of ssDNA aptamers (Sal 1 and Sal 2), target organism *Salmonella paratyphi A*, along with other related bacterial cultures like *Shigella flexnerri, Klebsiella pneumonia, Escherichia coli, Staphylococcus aureus, Pseudomonas aeruginosa* was coated onto a microtiter plate at 10^5^ cfu.mL^-1^ and ELASA was carried out and PBS kept as negative control.

### 2.5 Synthesis and characterization of Nanomaterials

#### 2.5.1 Synthesis and conjugation of quantum dot

The CdTe QDs were prepared as described in our previous reports (34) wherein 1 mM of CdCl_2_.5H2O along with 0.3 mM of TGA, 0.24 mM of Na_2_TeO_3_ and 0.1g of trisodium citrate, respectively was dissolved in 15 mL of ultra-pure water. The pH was adjusted to 10.5 and the mixture was subjected to vigorous shaking at ambient atmospheric condition. Upon coloration, the reaction was refluxed at 100° C in order to facilitate the growth of TGA capped quantum dots. Further, surface modified QDs were streptavidin functionalized by incubating with 200 μl of EDC and NHS at a ratio of 1:4 and allowed for constant stirring for 2 hours. The streptavidin functionalized QDs, biotin-labeled aptamers were anchored through avidin-biotin affinity (1:5; QD: DNA). The un-reacted fractions were removed by centrifugation at 13,000 rpm for 30 minutes, followed by subsequent washing of the pellets with ultra-pure water. These bioconjugated QDs were re-suspended in borate buffer (pH 8.5). The zeta potential and particle size distribution of CdTeQDs and bio-conjugated CdTeQDs were analyzed by differential light scattering (Malvern Zetasizer, US).

#### 2.5.2. Graphene oxide synthesis

Graphene Oxide (GO) was synthesized by Hummer’s method with modifications. Briefly, 3g of graphite powder was added to a mixture of concentrated sulphuric acid (60mL) and potassium permanganate (5g) and stirred at 25 °C for 120h. The potassium permanganate mediated oxidation was stopped with 100 mL of ice-cold deionized water and 5 mL of 30% H_2_O_2_. The resultant solution was filtered and washed with 1M HCl solution, and subsequently dialyzed (3.5kDa cut off membrane) against deionized water for 72h. The obtained GO was subjected to ultrasonication at 300 W, 25 % amplitude for 90 min and was lyophilized for further use. The morphological characterization was carried out with the Transmission electron microscope (JEM 2010) and Scanning Electron Microscope (FEI Quantasem). The powder X-ray diffraction analysis (Malvern panalytical) was performed with Cu monochromatized Kα radiation (λ = 1.541 Å) with an accelerating voltage of 40 kV. Thermal stability was analyzed with aid of Thermal Gravimetric Analysis.Surface area analysis was carried out with BET (Belsorp Mini, MicrotracBEL corp., Japan).

### 2.6 Sensor preparation

Graphene oxide prepared was dissolved in Milli-Q water and sonicated for 10 min and homogenous dispersion thus obtained were employed for sensor development.75nM aptamer labelled with QDs were prepared by diluting the stock in Phosphate buffer.

### 2.7 Development of GO/CdTeQDs-aptamer ensemble for *Salmonella paratyphi A* detection

A GO/CdTeQD-aptamer-based FRET assay was developed for rapid and sensitive detection of *Salmonella paratyphi A*. The QD-aptamer (75 nM) solution (in PBS; pH 7.4) was incubated with varying concentration of GO (0.01mg/mL to 1 mg/mL) and allowed for constant stirring for ~15 min. The fluorescence quenching was recorded with PL and optimum GO concentration was determined. The GO-Apt/QD complex was incubated under different *Salmonella paratyphi A* concentrations (10 cfu.mL^-1^ to 10^7^ cfu.mL^-1^) to assess the assay sensitivity. Similarly, the assay specificity was analyzed against different bacterial cultures (10^5^cfu.mL^-1^) such as *Salmonella paratyphi, Salmonella typhimurium*, and *Shigella flexnerii, E.coliO157:H7* and *Yersinia entreocolitica* cells. The fluorescent intensity values were recorded using Shimadzu RF 5301; Spectrofluorometer and Biotek Synergy H1 Hybrid multi-mode reader.

### 2.8 Evaluation of the GO/CdTeQD-aptamer-based FRET assay

The developed FRET assay was evaluated onto various artificially contaminated milk, meat and chicken samples to ascertain its robustness and feasibility.Different food matrices used for spiking studies were pre-evaluated for the presence of *Salmonella paratyphi A*. The meat and chicken samples (1g) were finely grounded and dissolved in PBS. The liquid carcass fraction was removed by gentle rocking for 2 hours and spiked with *Salmonella paratyphiA* (10^2^ cfu.mL^-1^to 10^4^ cfu.mL^-1^). Similarly, milk samples were directly spiked with *Salmonella paratyphiA* and diluted in PBS (1:10). The filtrate obtained was utilized for FRET-basedaptasensor assay, and their corresponding recovery percentage from each matrix was determined. The matrix interference of the developed assay was evaluated followed by measurement of inter and intra assay variation.The results thus obtained were compared with commercially available standard ELISA kit (MBS568055, My biosource, USA). Recovery percentage calculations are carried out as follows

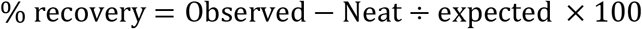

Wherein,

Observed – Obtained Spiked sample value,

Neat-Unspiked sample value,

Expected-Amount spiked into sample

### 2.9 CdTe-Graphene oxide interaction-Density functional theory (DFT) studies

The interaction of graphene oxide with CdTe quantum dot was studied with DFT employed in gaussian 09 packages^37^. The B3LYP level theory withLANL2DZ basis set was used to calculate the interaction energy and structural parameters. The TD-DFT studies were employed to calculate the absorption and emission wavelengths upon the interaction of QD and GO. The interaction energy had been corrected for basis set superposition error (BSSE) by applying counterpoise (CP) correction method of Boys and Bernard have given by

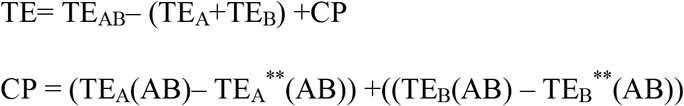

Where TE_A_(AB) and TE_B_(AB) represent the energy of quantum dot and graphene oxide the complex geometries and the terms TE_A_**(AB), and TE_B_**(AB) are the energies in the presence of respective ghost orbitals.

### 2.10 Aptamer-DNA gyrase interaction-Molecular dynamics simulation

The linear chain of the aptamer was constructed using YASARA structure package and subjected to an MD simulation in water to obtain the initial conformation. The initial conformation was further simulated for an indefinite time to obtain a stable 3D conformation. Further, the modelling of the ssDNA was continued in GROMACS 2018.1 ^39^ with CHARMM36 forcefield and TIP3P water model. The van der Waals interaction cut off distance of 1.2 nm with switching distance of 1 nm was used along with particle mesh Ewald method. The conformation was minimization using steepest descent method and equilibrated in NVT and NPT ensemble for 10 ns time. The production run of 500 ns was performed in NPT ensemble and trajectories were saved in each 100 ps interval. The temperature 310 K and pressure of 1 bar was maintained during the simulation using Langevin piston algorithm method. The periodic boundary conditions were applied in all three direction and trajectories were analysed using built tools in the GROMACS package. All the visualization was performed using PyMOL and Chimera software packages ^40^.

### 2.11 Statistical analysis

To ensure the consistency of the developed assay, all experiments were repeated thrice with a similar condition. The results obtained were represented as mean values with standard deviation. The significance of the data generated was determined using univariate (ANOVA) and student’s ‘T’ test. Results obtained were plotted using Graph Pad Prism 6.

## 3. Result and Discussion

The ssDNA aptamer generated against *Salmonella paratyphi* A by whole-cell SELEX method was characterized using biochemical assays. The high affinity and specificity of these aptamers having potential applications such as targeted drug delivery, biomarkers, and sensors. The generated aptamers are conjugated with quantum dot and graphene oxide and developed the potent sensor for the detection of *Salmonella paratyphi* A. Further, quantum chemical as well as molecular dynamics simulations were employed to understand the basic mechanism of sensor and action on the target protein.

### 3.1 Aptamer selection, interaction, and evaluation

Target specific aptamers against *Salmonella paratyphi* A were generated from a random sequence pool of oligonucleotides library by whole-cell SELEX method. The non-specific nucleotides that bind to close related pathogens of the *Salmonella* species were removed by seven rounds of definite target selection followed by four rounds of counter selection. Aptamer pool harvested after 11 rounds of selection remained stable with enhanced specificity towards target as monitored through Enzyme-Linked Aptamer Sandwich Assay (ELASA). Selected aptamer clone were subjected to sequencing and its corresponding secondary structure confirmations were predicted using M-fold software **(Figure.1)**. Structural stability of these clones was validated with aid of QGRS software, which evaluates clones on basis of their ability to form quadruplex structures and its corresponding G-score were tabulated in **Supplementary table 1**

**Fig 1:**
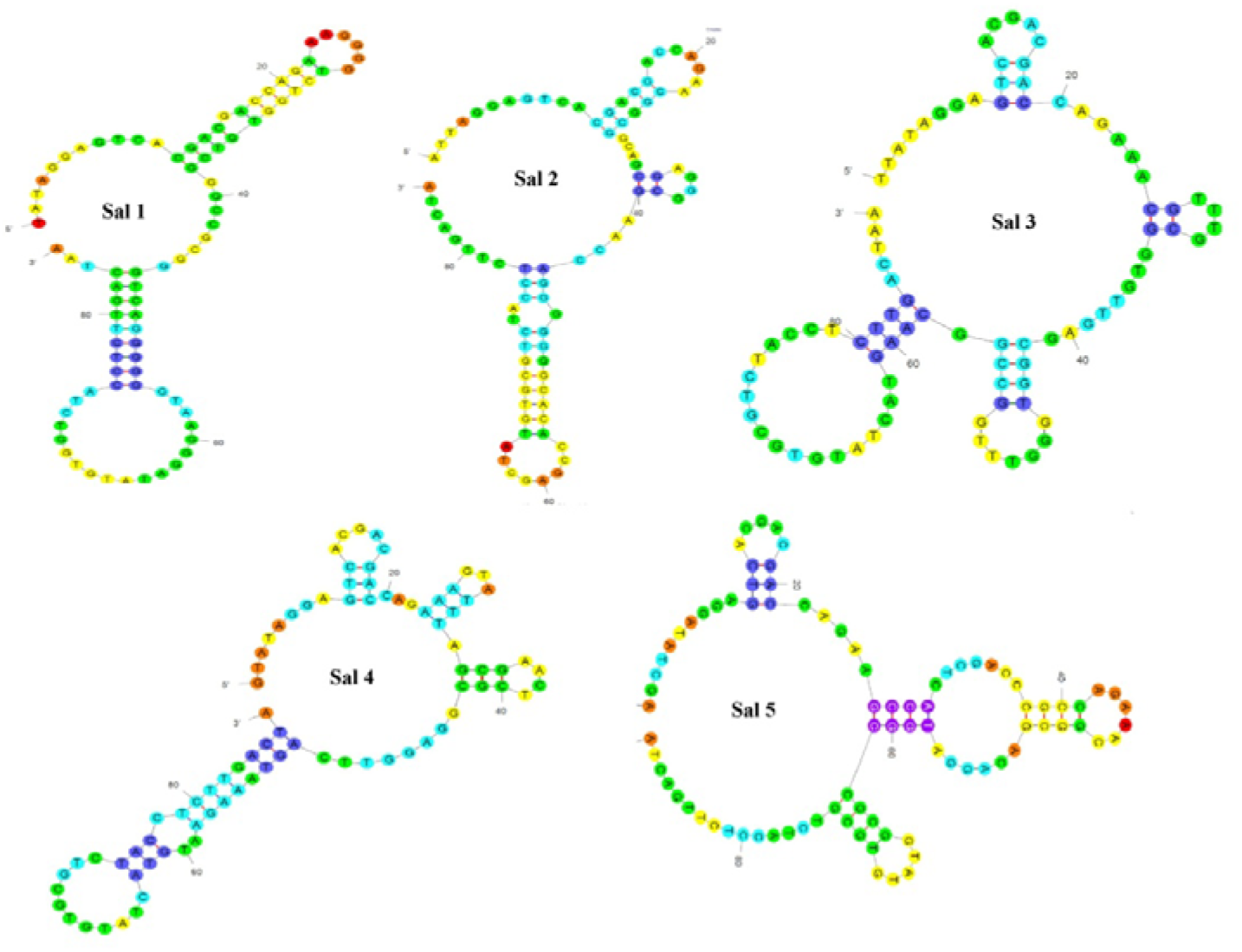
Secondary structures of generated aptamers. Secondary structures of aptamers generated towards *Salmonella paratyphi A* using M-fold software.

### 3.2 Characterization of generated aptamers by indirect ELASA

Two aptamers Sal1 and Sal2 pose a limit of detection of 10 and 10^3^ cfu.mL^-1^, respectively **(Figure 2a)**. As seen from **Figure 2b**, Sal1 and Sal2 show high affinity towards *Salmonella paratyphiA* without any cross-reactivity that underlie aptamers specificity towards target pathogen. Titer value estimation showed reactivity up to 1:51,200 corresponding to the concentration of 120 ng /mL (**Figure 2c**). Based on its characterization data Sal1 aptamer was selected for the design of FRET-based aptasensor platform towards *Salmonella paratyphi A*.

**Figure 2:**
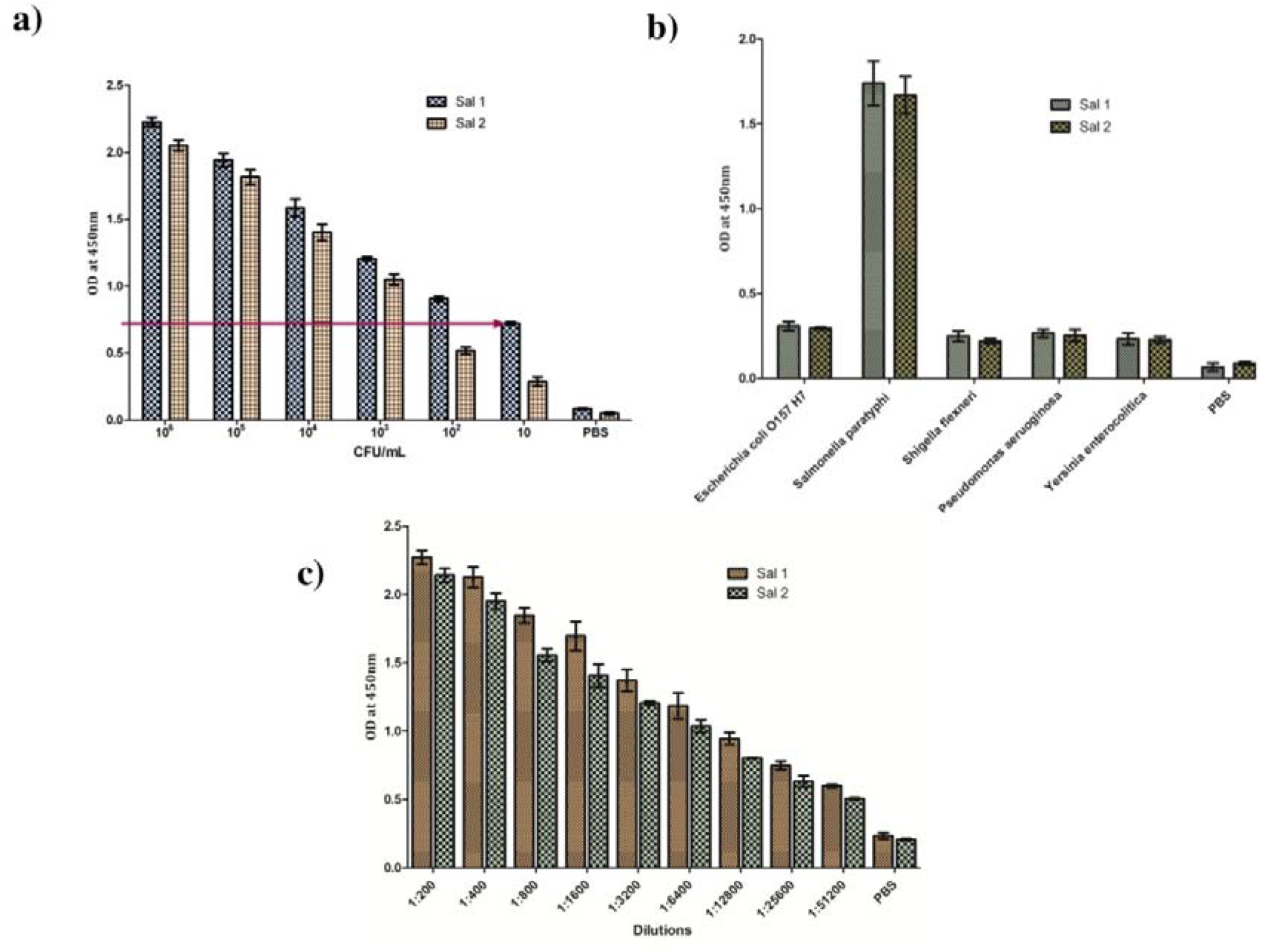
Characterization of *Salmonella paratyphi A* (Sal1 and Sal2) aptamers by Indirect ELASA. **a**) Sensitivity determination of Sal1 and Sal2 aptamers against target pathogen. **b**) Specificity analysis of aptamers against 1) PBS, 2) *E.coli* O157:H7,3) *Salmonella paratyphi A*, 4) *Shigella flexneri* 5) *Salmonella typhimurium*, 6) *Yersinia enterocolitica*, **c)** Titre value estimation of Sal1 and Sal2 aptamers.

### 3.3 Synthesis and characterization of nanomaterials

#### 3.3.1 Synthesis and characterization of Graphene Oxide

Graphene oxide was synthesized through Hummer’s method and characterized using XRD, TGA, SEM, and TEM. TEM analysis revealed wrinkled and transparent sheet signifying the presence of epoxy groups on the surface (**Figure 3a**). Also, the SEM micrograph depicted a typical rippled structure sealed throughout the functionalized surface (**Figure 3b**) ^40, 41^. XRD shows characterized peak for graphene and graphene oxide and at 2θ = 26.6 and 10.5 °with interlayer spacing of 0.85 nm (Figure). Further, the broader peak of GO signifies regular graphite crystalline structure damage by oxidation (**Figure 3c**). The thermogravimetric analysis revealed the time dependant weight loss of the GO as shown in **Figure 3d**. An initial loss of 10% at 100° C increased significantly upon higher temperatures as a result of interlamellar water loss and decomposition of labile oxygen-containing groups such as epoxy, carboxyl and hydroxyl ^42–44^. Furthermore, the surface area and pore volume computed by Brunauer-Emmett-Teller (BET) method for graphene and GO is given **Supplementary Table 2**, a slight variation is due to fractional restacking of graphenelayers with locally blocked pores^45–47^.

**Figure 3:**
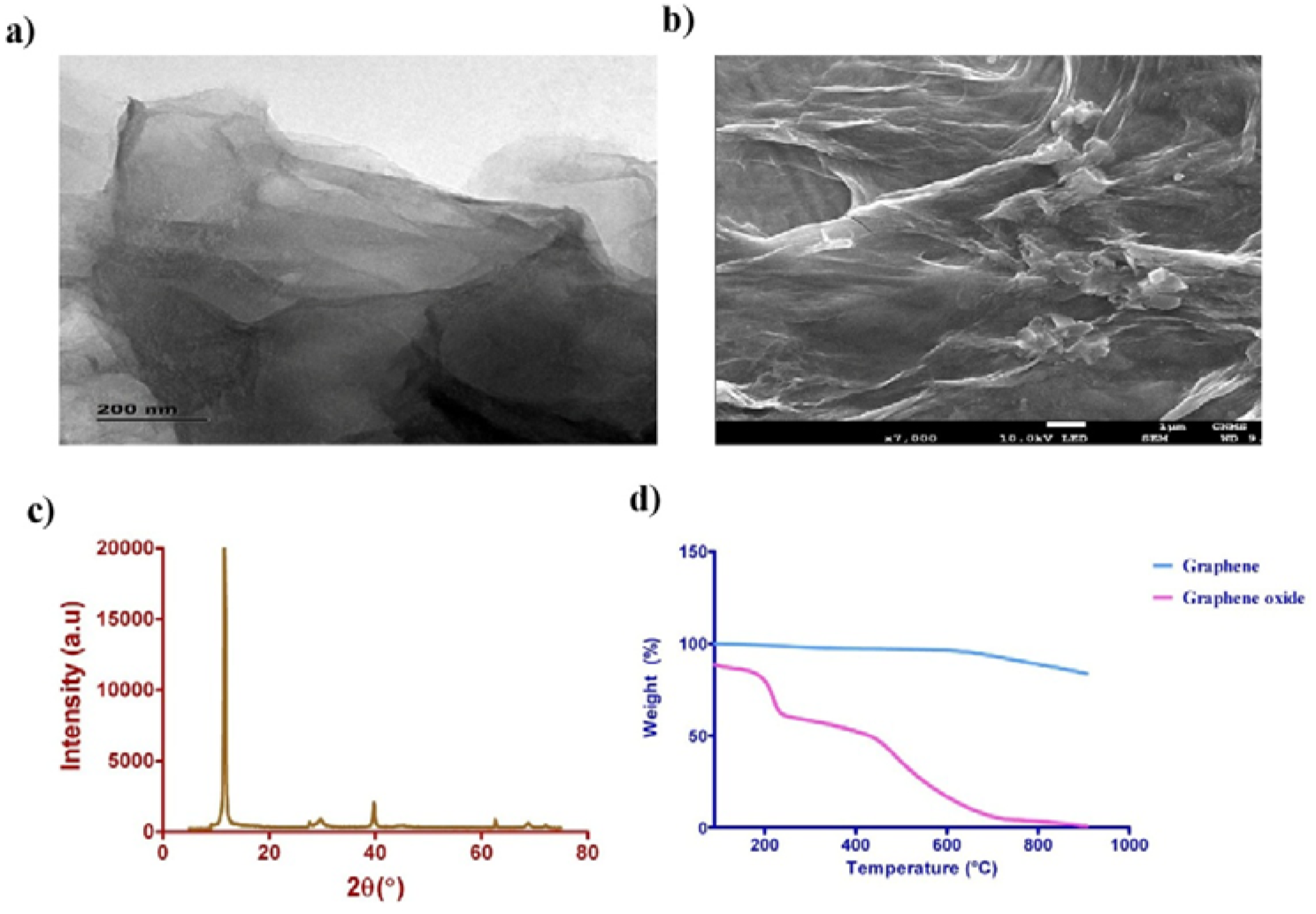
Synthesised Graphene Oxide characterization. **a)** TEM and SEM characterization of Graphene Oxide, **b)** X-ray diffraction pattern of synthesised GO, **c)** Thermo gravimetric analysis of graphene oxide.

#### 3.3.2 Characterization of CdTe quantum dot

The quantum dot exhibits a broad range of absorption without any donor and acceptor spectrum overlap. As a result, QDs are preferred as an ideal candidate for FRET-based assays^48^. Bio-labelling of streptavidin functionalized QDs with biotin-tagged aptamers were facilitated through avidin-biotin interaction ^49^ The UV Vis spectrum of Apt/QDs complex revealed peak broadening associated with Sal 1 aptamer conjugation onto the nanoparticle surface (**Figure 4a**). A decrease in PL intensity upon bio-conjugation of QDs reveals their energy transfer towards aptamers (**Figure 4b**).

**Figure 4:**
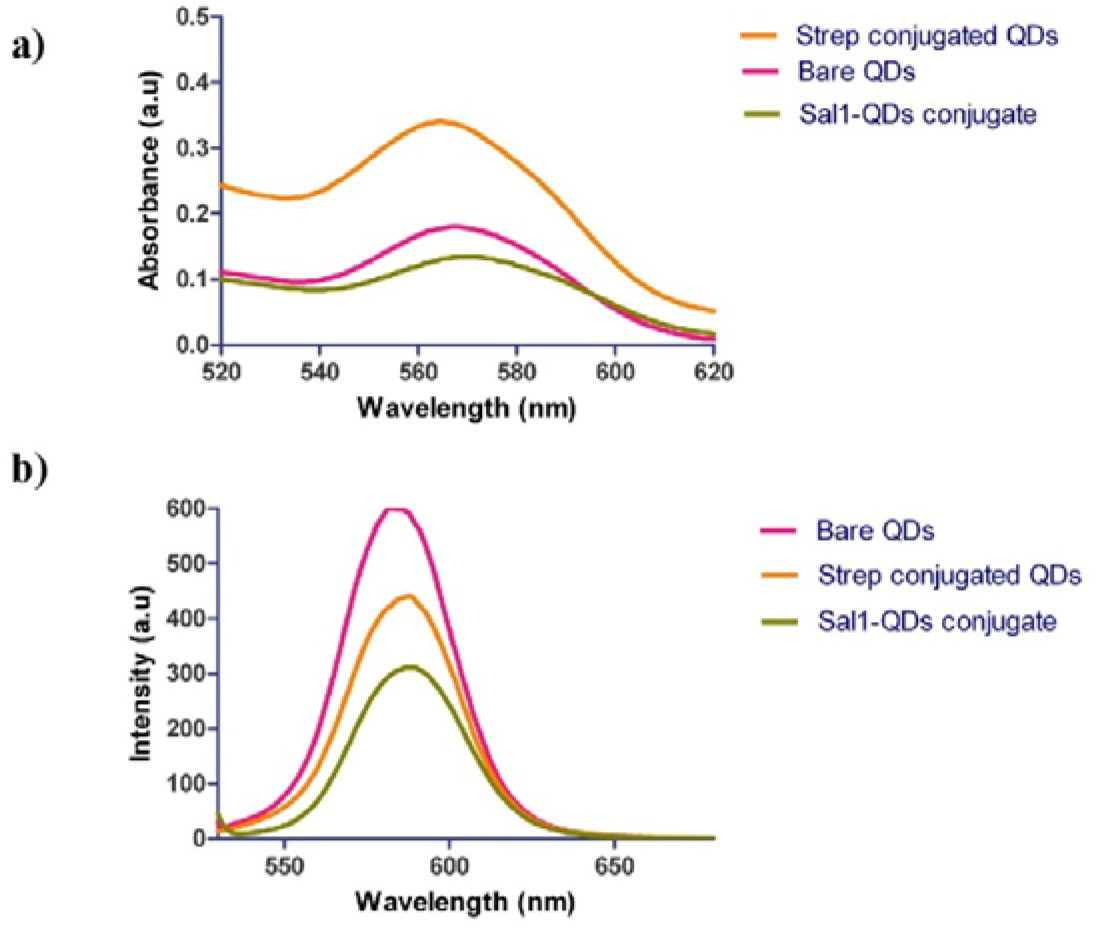
Characterization of Aptamer –QD conjugate. **a)** UV-visible spectra characterization of surface modified CdTe QDs, **b)** PL spectra characterization of surface modified CdTe QDs and conjugate, **c)** DLS characterization of synthesized and functionalized QDs.

### 3.4 GO/CdTeQDs-aptamer ensemble based FRET assay for *Salmonella paratyphi A* detection

The platform is based on turn on/off fluorescence between an electron donor and acceptor pairs. The system consists of CdTe quantum dot conjugated aptamer as the donor and GO as acceptor. The ssDNA aptamers adsorbed onto the GO surface and adsorption can be linked to hydrophobic and π-stacking interaction of the fluorescent nanomaterial as well as the interactions due to the nucleobases with the aromatic region of the GO. This interaction considerably quenches the fluorescence of aptamer linked QDs emphasizing an excellent electronic transference and conductivity of graphene ^50^. The FRET-based quenching is reversed in the presence of target. The gradual increase in fluorescence amid the target presence indicates ssDNA desorption from the GO surface. Therefore, the features such as adsorption-desorption and fluorescence quenching ability with a wide range of fluorophores makes GO as the prominent material of choice for biosensor development.

### 3.5 CdTe-Graphene oxide interaction-Density functional theory-based calculations

The Cd_6_Te_6_ quantum dot (QD) and graphene oxide conjugated quantum dot ground state and excited state properties were studied using density functional theory and time-dependent density function theory methods (TD-DFT) as implemented in the Gaussian 09 software package. Random location was considered such that the graphene oxide would bind/interact strongly with quantum dot. The geometry optimization and absorption-emission spectra wereperformed with B3LYP functional and LANL2D basis set. The HOMO-LUMO energy gap calculated as E=HOMO-LUMO, where HOMO and LUMO referred to the highest occupied and lowest unoccupied molecular orbitals, respectively. The change in HOMO, LUMO (**Figure 5**) and the corresponding transition were observed before and after the binding of the graphene oxide as **(Supplementary Figure 1)**. The TD-DFT method provided the excited state properties of the bare quantum dot and graphene oxide conjugated quantum dot in the gas phase, the peaks of the absorption spectra of the quantum dot and conjugated structure is given in **Figure 6**. Absorption spectra of QD show peaks at 394 nm, 428 nm and 480 nm for corresponding HOMO→LUMO transition and the energy values have been limited to 7.0 eV and excitation up to 12 states. The excitations have low oscillator strength because of less overlap between conjugated graphene and quantum dots. The transition occurs from HOMO (1) to LUMO with an energy difference of 3.05 eV (480 nm) and HOMO (5) →LUMO as shown in the MO diagram **(Supplementary Figure 1)**. The conjugated structure shows absorption peaks at 777 nm,785 nm,806 nm,881 nm,963 nm,1035 nm,1310 nm,1449 nm and 1837 nm corresponding to the transition from α-α orbitals and β-βorbitals with an energy difference of HOMO-LUMO as 0.66 eV, 0.86 eV, 0.95 eV, 1.2 eV, 1.23 eV, 1.4 eV, 1.46 eV, 1.51 eV, 1.53 eV 1.57 eV and 1.59 eV. The abortion and emission spectra included several excited states, where the absorption peaks range from 200 nm to 300 nm and the emission has a wider range from 300 to 700 nm. The higher excited state is responsible for the several strong peaks in the range of shorter wavelength. The quantum dot spectra show a small redshift, whereas graphene oxide conjugated QD shows a blue shift with induced changes on the quantum dot structure by graphene oxide (**Figure 6**).Further, the natural transition orbital analysis (NTO) for the lowest excitonic transition of the graphene oxide conjugated quantum dots were calculated and given in **Figure 7**. In the excitonic excited states, the photogenerated hole of CdTe quantum dot resides mainly on Te and electron residues primarily in Cd, electron distribution was seen as spherical like density distribution with a maximum at the center. The NTO surfaces for the 12 states of the bare quantum dot do not show any significant changes and the electron density shows thick spherical shape around individual atoms. The distribution of photogenerated hole and particle are found to be located around Te atoms at the first excited states, and several s-shaped spherical distributions around most of the atoms of the QD at the 2nd excited states were observed. Observing all the 12 states of the conjugated quantum dots shows the possibilities of the energy transition. These results implicate that the interaction of grapheneled to the modification of electron wave function in the quantum dot, which further reveals the energy transfer as observed in the experimental results.

**Figure 5.**
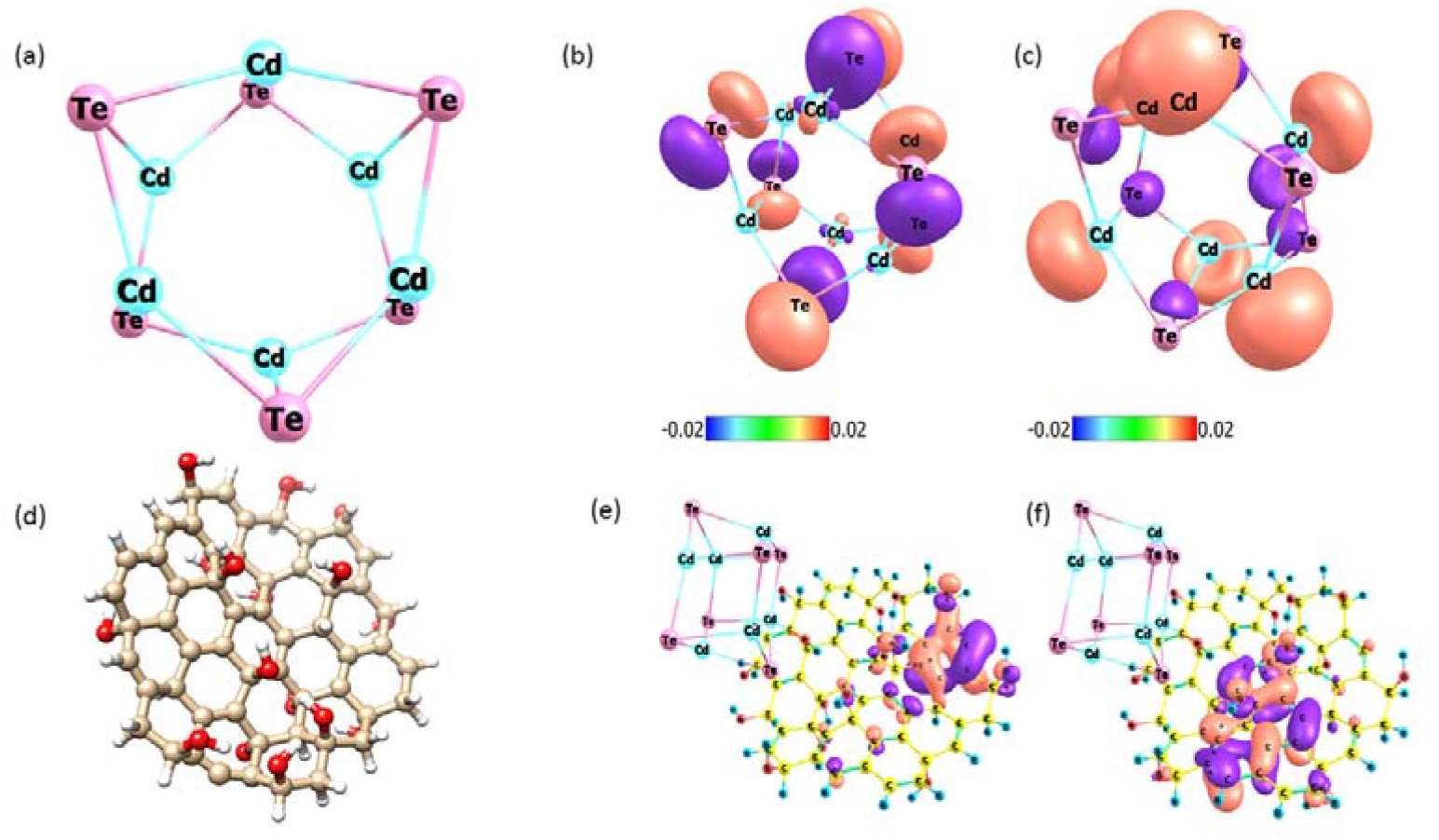
(a)-(d) represent the optimized three-dimensional structure of CdTe quantum dot and graphene oxide. (b) Graphical representation of HOMO, LUMO state of bare quantum dot and (e) represent the HOMO, LUMO shift when interacting with graphene layer

**Figure 6.**
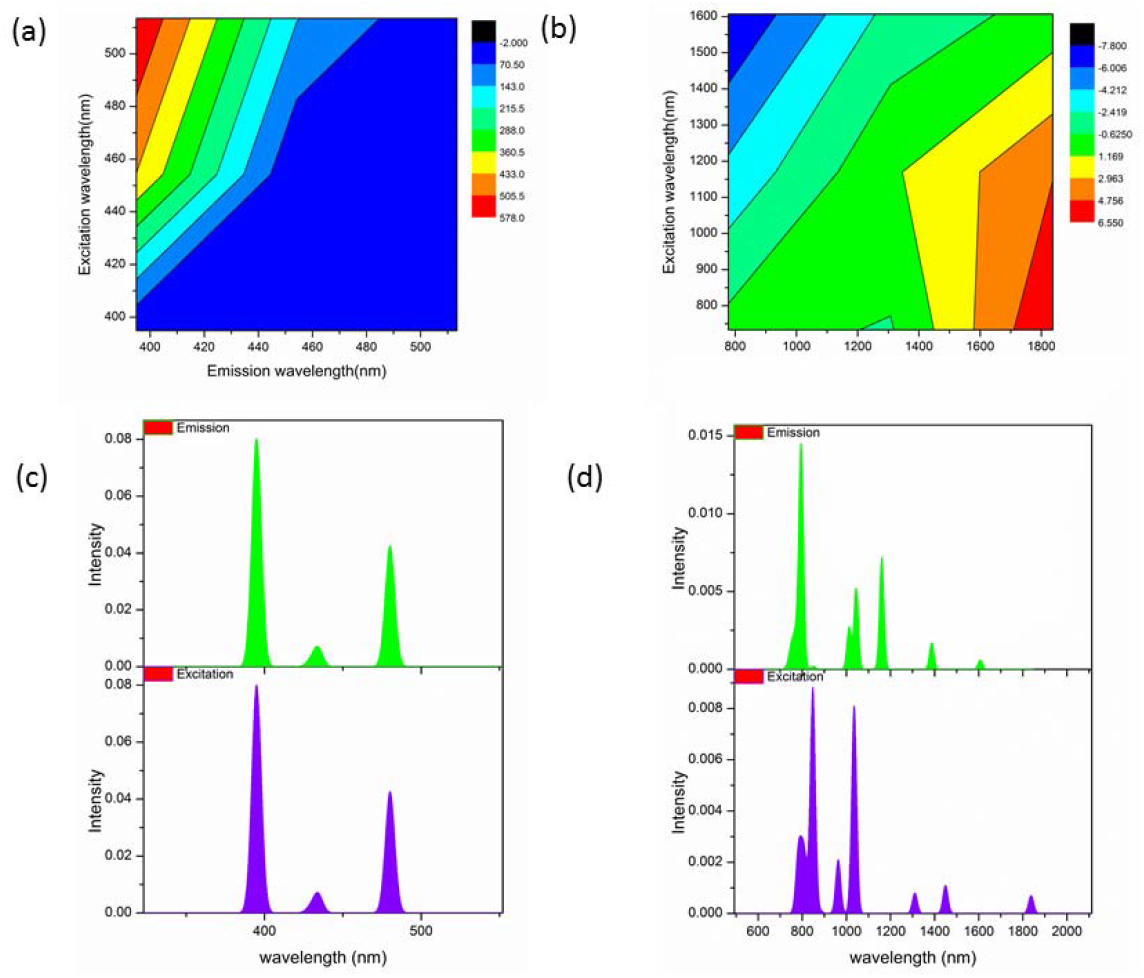
(a)-(b) The fluorescent shift before and after binding of quantum dot with graphene oxide. The corresponding absorption and emission spectra are depicted in (c) and (d).

**Figure 7.**
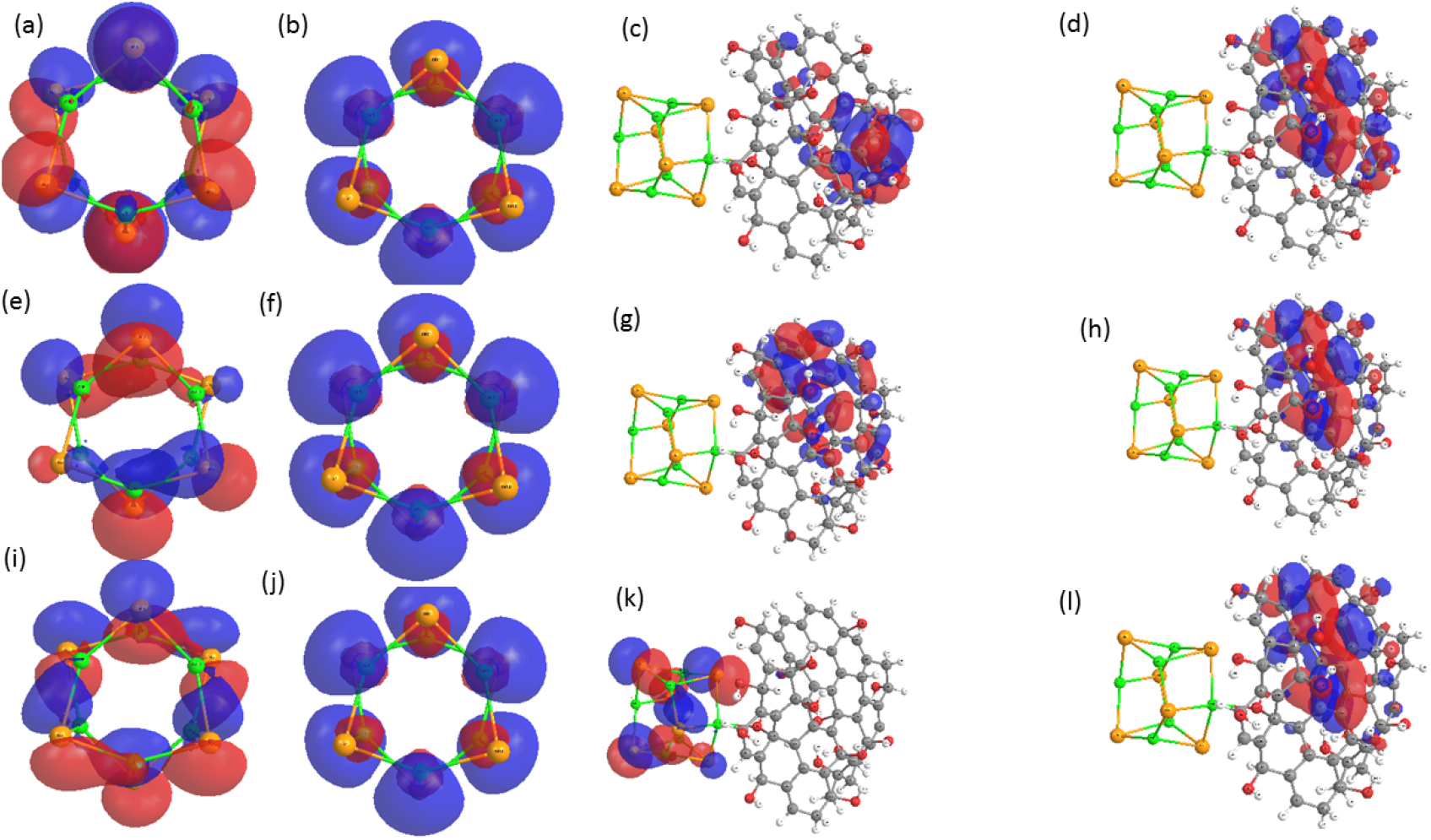
Represent the natural transition orbital analysis (NTO) for the lowest excitonic transition of the quantum dot and graphene oxide conjugated quantum dots. (a)-(d) represent at the first excitation state (e)-(h) represent at sixth excitation state and (i)-(l) represent for 12^th^excitation state.

### 3.6 Optimization of GO concentration for the sensor

GO at higher concentrations permits the adsorption of reaction components onto its surface whereas lower levels being incapable of the same action makes it an efficient quencher molecule. Hence, the concentration of GO below which no remarkable quenching is observed is selected as the optimum for sensor development. The fluorescence intensity of QDs labelled aptamer at a varying concentration of GO is demonstrated in **Figure 8**. A decrease in the intensity can be observed with a gradual increase in GO concentration. The complete quenching attained at 0.1 mg/mL chosen as the optimum for the FRET-based platform development. The energy transfer between QDs and GO facilitates fluorescence quenching due to the π rich electronic structure along with its phenomenal π-π stacking. Furthermore, an enhanced quenching observed in the present scenario can be attributed to non-covalent interaction of aptamers with GO.

**Figure 8:**
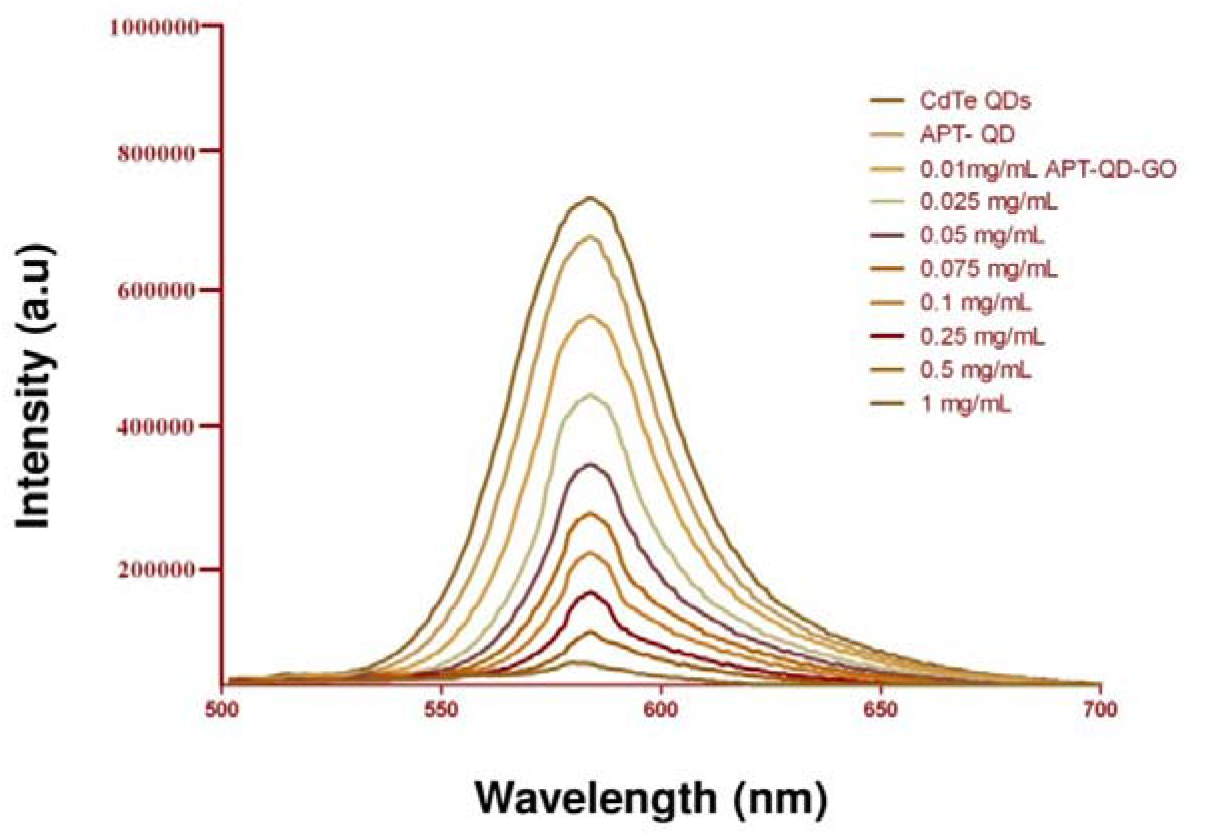
Optimization of GO concentration for FRET. Varying concentration of GO 0.01, 0.025, 0.05, 0.075, 0.1, 0.25, 0.5, 1 mg /mL were utilized to study the quenching efficacy of synthesised GO.

### 3.7 Specificity and sensitivity of FRET aptasensor

The selectivity of FRET-based aptasensor assessed by employing different non-specific bacterial targets such as *Escherichia coli* O157:H7, *Shigella Flexneri, Yersinia enterocolitica*, and *Salmonella typhimurium*. Their cross-reactivity analysis showed remarkable specificity towards *Salmonella paratyphiA* and none of the other examined organisms exhibited any fluorescence intensity (**Figure 9**). Therefore, this validates the high selectivity of the developed aptamer-based FRET biosensor for *Salmonella paratyphi A* detection. Similarly, their sensitivity evaluations at various cell densities (10 to 10^7^ cfu.mL-1) carried out to determine the limit of detection (**Figure 10a**). Under the defined experimental condition, enhancement of fluorescence intensity upon correspondingtarget concentrations indicated a good linear relationship both with an R^2^ value of 0.9591 (**Figure 10b**). The estimated limit of detection (LOD) for the developed technique showed as low as 10cfu.mL^-1.^This probe works directly by mixing the components aptamer-QDs, GO and target pathogen and its sensitivity range are quite promising compared to methods reported earlier **(Table 2)**.

**Figure 9:**
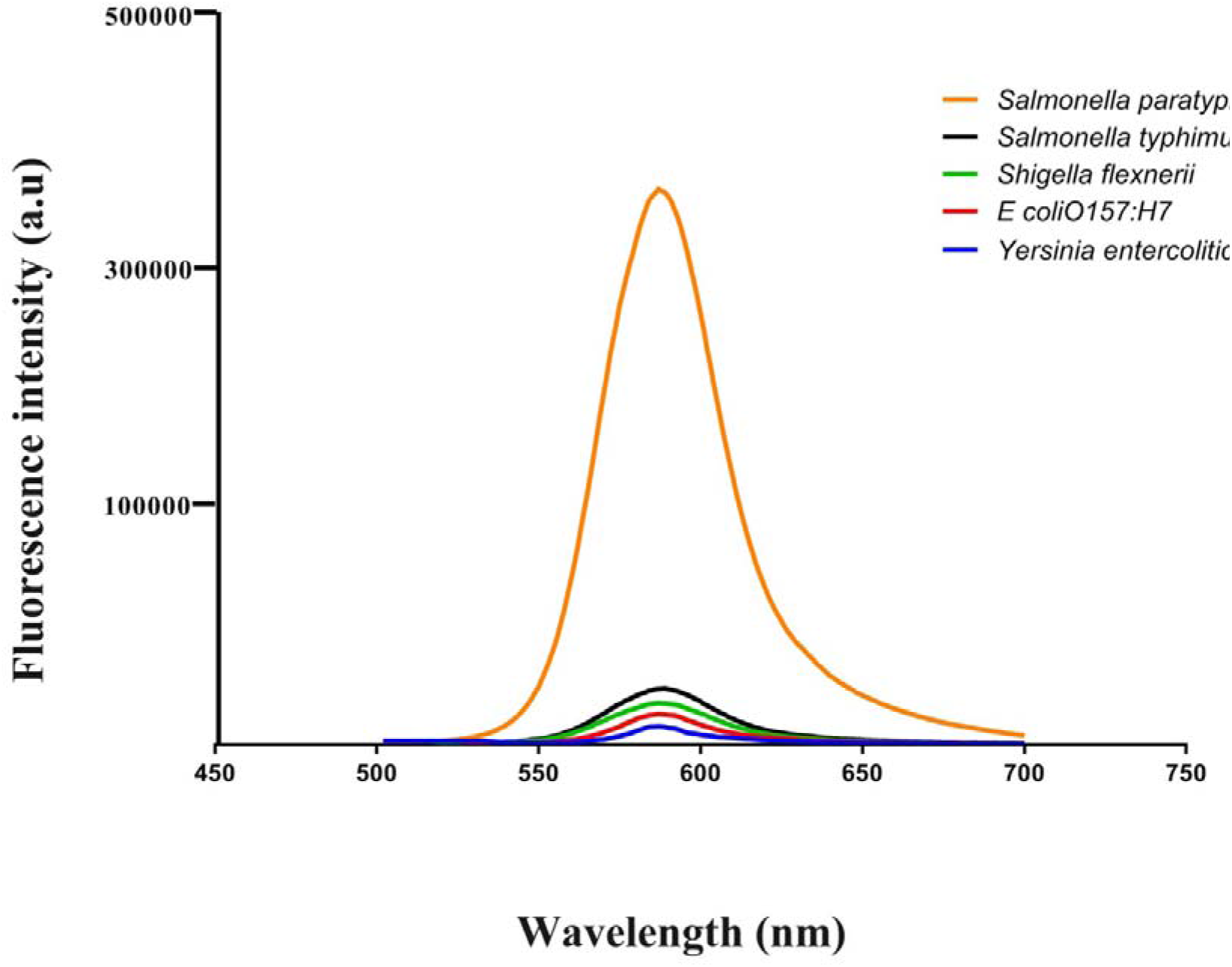
Specificity evaluation of developed FRET. Specificity of developed FRET was verified by recording fluorescence with various bacterial targets.

**Figure 10:**
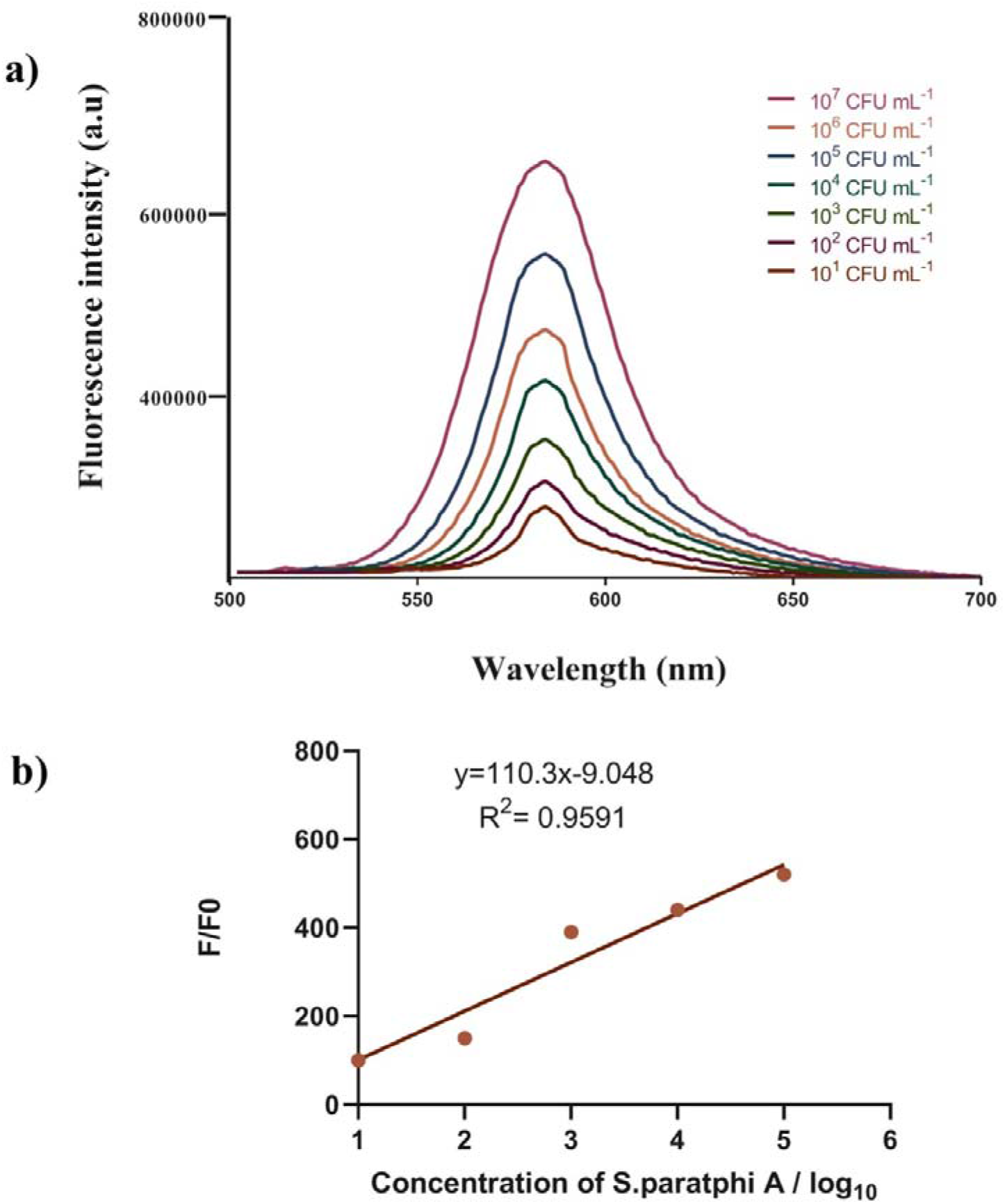
Sensitivity evaluation of FRET aptasensor. **a)** Sensitivity of designed platform was evaluated to determine its limit of detection against varying CFU/mL concentration of *Salmonella paratyphi* A 10^7^ CFU/mL to 10^1^ CFU/mL. **b)** Calibration plot of fluorescence intensity against varying *Salmonella paratyphi* A concentration.

**Table 2:** Assessment of developed aptasensor onto standard cultures and field samples.

| **SL. No** | **Organism** | **Source** | **Developed FRET based aptasensor** | **Standard ELISA(MBS568055)** |
| --- | --- | --- | --- | --- |
| 1 | *Salmonella paratyphi A* | ATCC 9150 | + | + |
| 2 | *Klebsiella pneumonia* | ATCC 12022 | - | - |
| 3 | *Salmonella typhimurium* | ATCC 14028 | - | - |
| 4 | *Pseudomonas aeruginosa* | ATCC 27853 | - | - |
| 5 | *Shigella sonnei* | ATCC 25931 | - | - |
| **Spiked Isolates** |  |  |  |  |
| 1 | Chicken isolate 01 | NA | + | + |
| 2 | Chicken isolate 02 | NA | + | + |
| 3 | Chicken isolate 03 | NA | + | + |
| 4 | Milk isolate 01 | NA | + | + |
| 5 | Milk isolate 02 | NA | + | + |
| 6 | Milk isolate 03 | NA | + | + |
| 7 | Meat isolate 01 | NA | + | + |
| 8 | Meat Isolate 02 | NA | + | + |
| 9 | Meat isolate 03 | NA | + | + |

### 3.8 Detection of *Salmonella paratyphi A* from artificially spiked samples

There is a high possibility for the food samples contaminated with *Salmonella paratyphiA* also contain other bacterial species. Though the above section validated specificity of the developed system, the present study evaluated different food matrices influence namely meat, chicken and milk samples. Their impact assessment carried out by spiking each matrix with *Salmonella paratyphiA* and estimating the total recovery percentage of the organisms. The same has been co evaluated with commercially available kit. There was no significant difference observed between the compared methods suggesting the feasibility of the assay for real-time food sample analysis. Likewise, no matrix interference effects recorded for the developed system as represented by Supplementary figure 2(SF 2).The intra assay coefficient of variation for the developed system found to be between 2.5% to 5.5% with recovery percentage of 102% to 80% (**Table 3**). The interassay coefficient variation of 2.9% to 6% with recovery of 104% to 83% highlights platform efficacy. The fluorescence thus recorded highlights the fluorescence recovery potential of the developed system in target presence. Thus, the precision and reproducibility of the developed platform were evident in the level of recovery percentage from food sample analysis. The developed FRET-based platform compared in terms of its sensitivity with reported platforms is listed in **Table 4**.

**Table 3:**
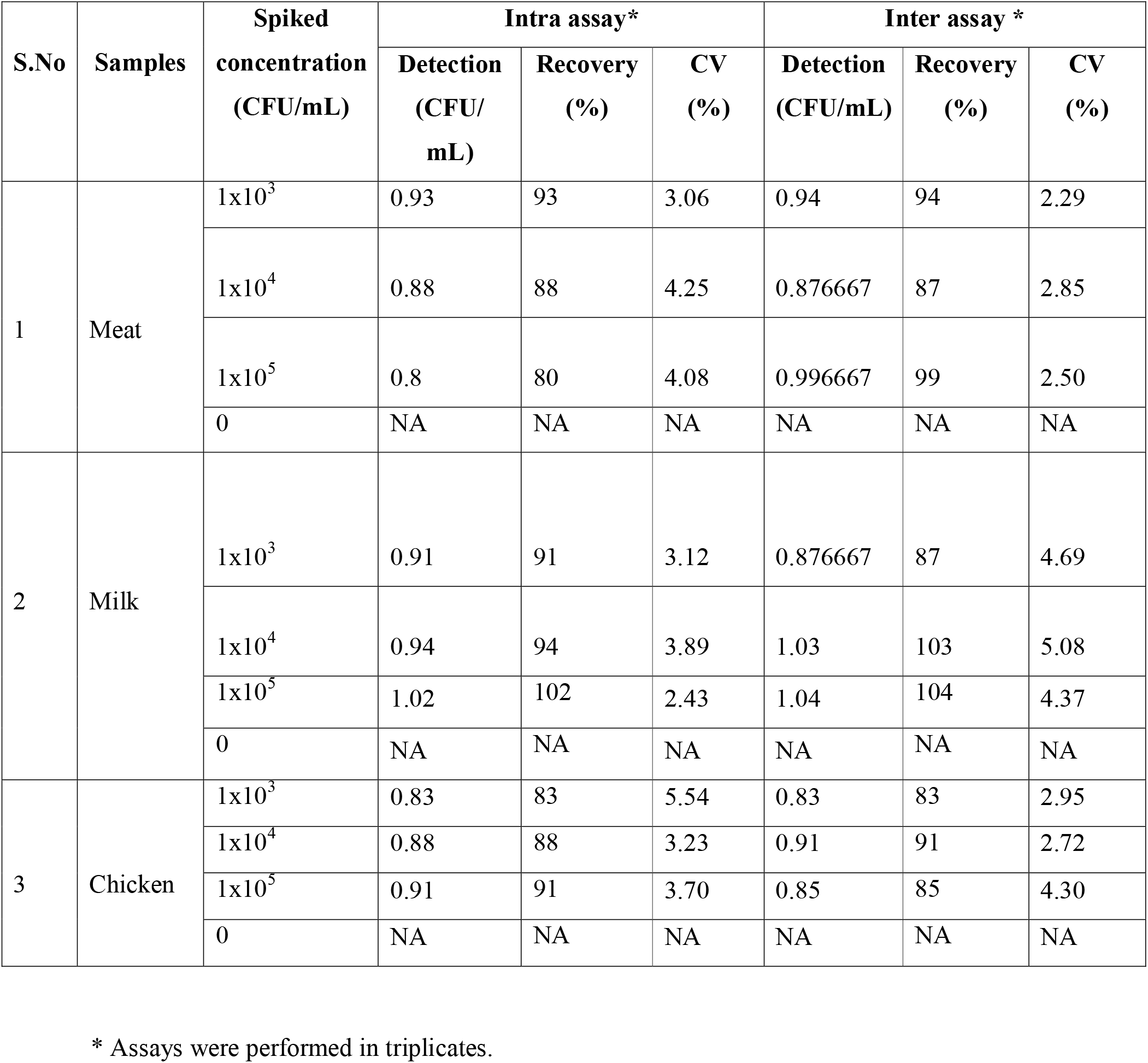
Recovery analysis from spiked samples using FRET based aptasensor.

**Table 4:** Reported sensor platforms for *Salmonella paratyphi A* detection.

| Pathogen | Detection method | Materials used | LOD | Reference |
| --- | --- | --- | --- | --- |
| *Salmonella paratyphi A* | Fluorescence | Single-walled carbon nanotubes (SWNTs) and DNAzyme-labeled aptamer detection probes | $10^4$ cfu/mL | <sup>53</sup> |
| | Fluorescence | DNAzyme and fluorescein isothiocyanate-labeled aptamers | $10^5$ cfu/mL | <sup>54</sup> |
| | PCR | Multiplex PCR | $10^2$ cfu/mL | <sup>55</sup> |
| | Fluorescence | DNase I mediated signal amplification with FAM labelled aptamers | $10^2$ cfu/mL | <sup>56</sup> |
|  | Fluorescence | GO-Aptamer labelled with QDs | 10 cfu/ mL | Present work |

### 3.9 Molecular modelling of ssDNA aptamer

The three-dimensional structure of ssDNA aptamer was modelled using molecular dynamics simulation. The linear chain of the ssDNA begins to fold at 50 ns and stable conformation obtained at the end of 450 ns simulation time. The 450 ns simulation was performed in three intervals and in each interval the simulation system was redefined to reduce the computational cost by reducing the number of water molecules, final simulation system contains 98,649 atoms. The ssDNA three-dimensional conformation evolves as a function of time and after 400 ns the ssDNA reaches stable conformation **(Supplementary Figure 3)** and with no further significant structural changes were observed. The root means square deviation of the backbone atoms and solvation energy in each simulation steps have been computed. The number of hydrogen bonds formed between each base pair has calculated and found that a minimum of 5-10 conventional hydrogen bonds was present during the simulation time **(Supplementary Figure 3)**. The hydrogen bonds and non-bonded interaction energies between these pairs help the ssDNA to reach a stable three-dimensional structure. The modelled ssDNA structure was used to perform a docking with DNA gyrase protein to elucidate the binding mechanism.

### 3.10 Interaction and binding of aptamer with DNA gyrase

We have considered DNA gyrase as a target protein for the modelled aptamer to elucidate the interaction and mechanism of inhibition of the protein. DNA gyrase is an essential bacterial enzyme that catalyses the ATP-dependent negative super-coiling of double-stranded closed-circular DNA. The gyrase is a prominent target for antibiotics because of its presence in prokaryotes and some eukaryote. The gyrase structure was modelled using ITASSER server ^51^ and 50 ns MD simulations were performed to remove the bad contacts and steric hindrances. The final confirmation of the gyrase was evaluated using PROCHECK tool ^52^. Further, gyrase-ssDNA complex structure was used to perform 100 ns MD simulations in aqueous media.The structural stability and convergence of the simulation were assessed by calculating backbone RMSD and Cα-RMSF of the protein in both apo and aptamer bound complexes. The apo form of gyrase shows high fluctuations in RMSD with an average of 1 nm, whereas the aptamer bound gyrase shows more stable RMSD values with an average of 0.5 nm as shown in **Figure 11**. RMSF calculation also depicts the fewer fluctuations of the gyrase after aptamer binding, which indicates the less conformational freedoms of gyrase to fluctuation due to the binding ofthe aptamer. Further, PCA analysis was performed to understand how gyrase conformational free energy landscape change upon binding of the aptamer.The PCA was based on Cα atoms of gyrase and it leads to a mathematical equivalence to quasi-harmonic analysis, which is used to identify low-frequency normal modes in proteins in the mass-weighted quasi-harmonic approximation. We have considered five principal components (PCs) based root means square deviations of Cα atoms and PC1 and radius of gyration were used to obtain the free energy surfaces. Free energy landscape plotted based on the PCs shows unbound gyrase having multiple peaks converged at various intervals. The aptamer bound landscape shows a broad and stable convergence of energy at two places. This result demonstrates that sampled conformation of the aptamer-bound gyrase can be found in the structural ensemble of the aptamer-unbound gyrase. Additionally, under the same amount of sampling, the gyrase without aptamer has more unique conformations which are not accessible after aptamer-binding **(Figure 12)**. This indicates that binding of aptamer significantly reduced the conformation freedom of the protein and it may reduce the functioning of the protein.

**Figure 11:**
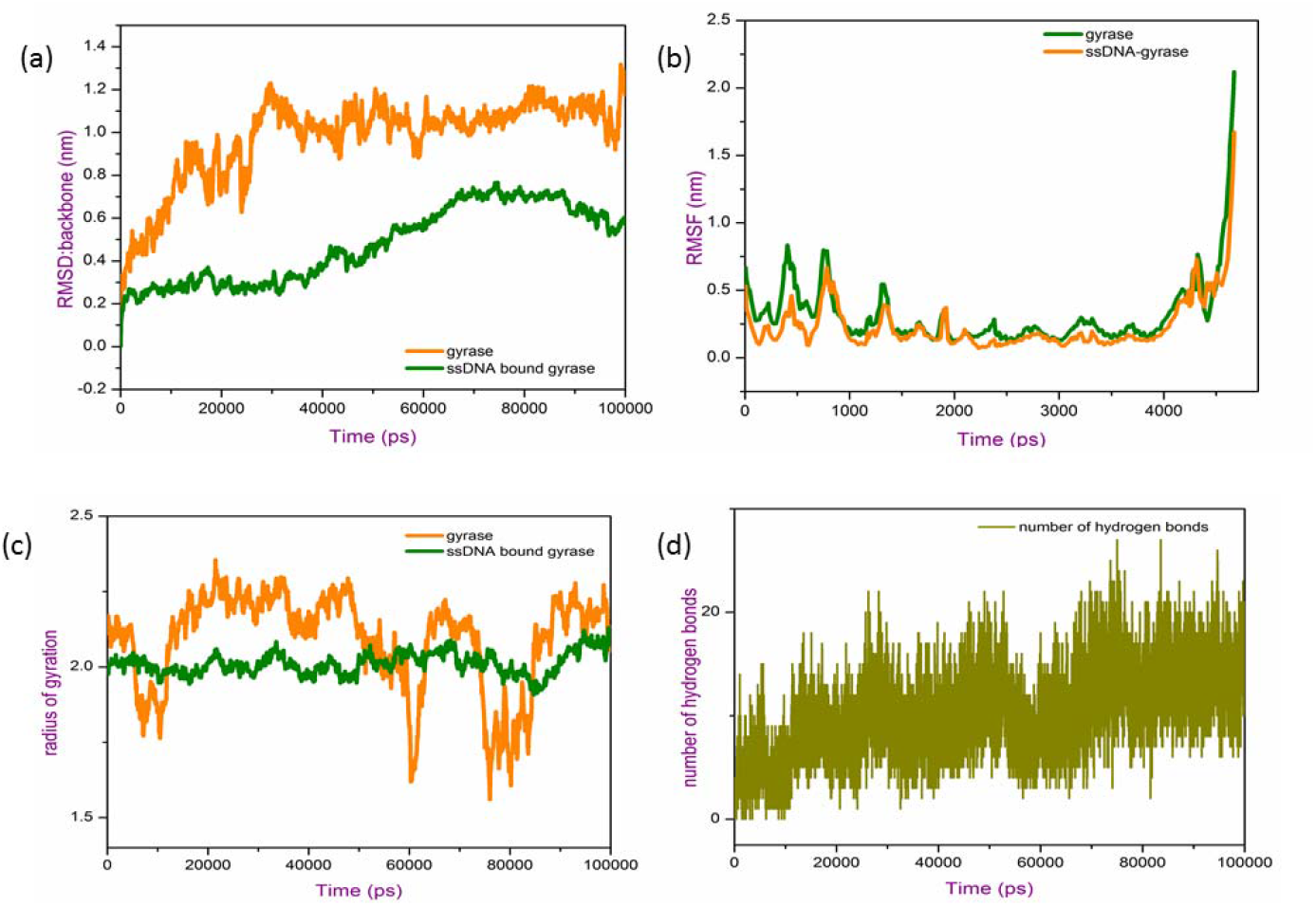
(a)Represent the root means square deviation of gyrase backbone atoms, native (orange) and ssDNA bound form (green). (b) represent the RMSF, (c) radius of gyration (d) number of hydrogen bond formed between ssDNA and gyrase protein.

**Figure 12:**
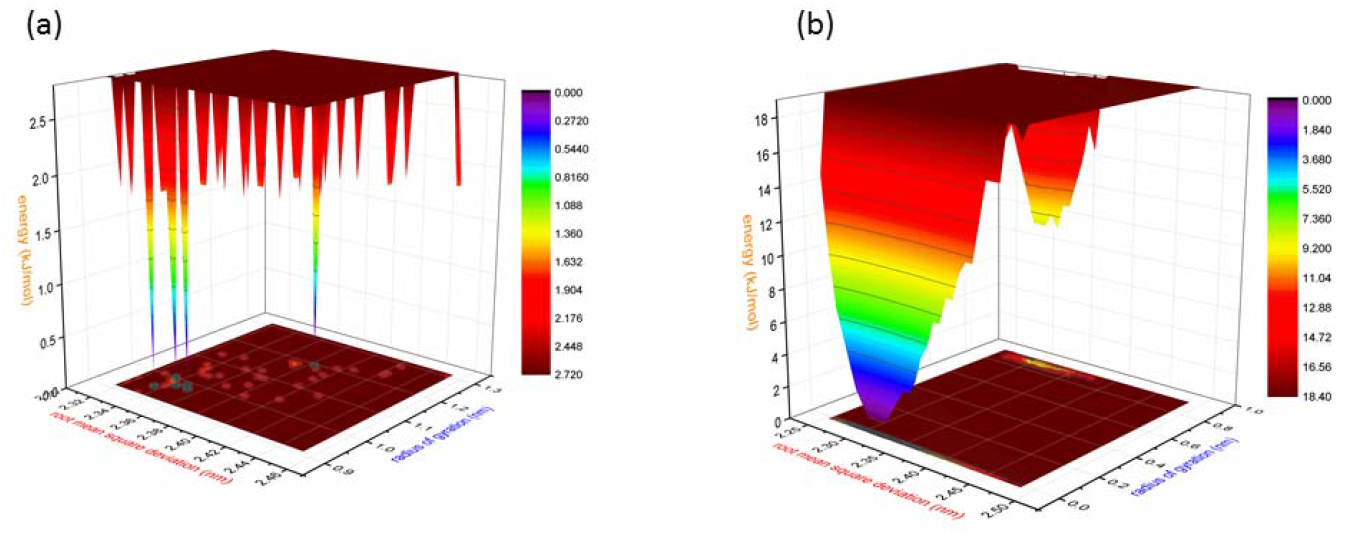
The Free energy landscape of ssDNA obtained for PC1 and radius of gyration (a) represent native protein while (b) represents ssDNA bounded gyrase.

The non-bonded interactions energies such as electrostatic and van der Waals interaction between the aptamer and gyrase protein have been calculated **(Supplementary Figure 4)**. The strong electrostatic and van der Waals energy between the aptamer and protein help to stabilize or freeze the protein from its functioning. Due to the long coiled and folded structure, the aptamer can access the large surface area of the protein. Further, the hydrogen bond between the aptamer and protein were calculated and it is found that an average of ten hydrogen bond was observed in each frame of the simulation. The high number of hydrogen bonds also depicts the strong affinity of the aptamer towards the gyrase. Furthermore, MM-PBSA calculation confirmed the strong affinity of aptamer molecules with gyrase protein. The electrostatic and van der Waal’s interaction energy was favouring for the binding of the aptamer and the summary of MMPBSA calculation is given in the **Table 5**.

**Table 5:** The contribution of different energy terms towards the total binding energy between ssDNA and gyrase protein by MM-PBSA method.

| **ssDNA-Gyrase binding** | **van der Waal energy (kJ/mol)** | **Electrostatic energy (kJ/mol)** | **Polar solvation energy (kJ/mol)** | **SASA energy (kJ/mol)** | **Total Binding energy (kJ/mol)** |
| --- | --- | --- | --- | --- | --- |
|  | -742.57 ± 3.4 | -2277.02 ± 4.5 | 2045.14 ± 2.04 | -56.86 ± 6.47 | -1031.31 ± 4.3 |

## Conclusion

We have developed a FRET-based aptasensor sensing platform for *Salmonella paratyphi A* detection. The assay includes distinct advantages like simple design, high specificity, and sensitivity with a limit of detection of 10cfu.mL^-1^. Prominent recovery percentage (87%-102%) and negligible matrix interference from the spiked food samples showed compatibility for routine screening of field samples. Significant fluorescence enhancement recorded upon the target GO-aptamer complex formation whereas other bacterial strains didn’t produce any similar effects. Thus, with a comprehensible design and highly efficient performance, the developed platform could be applicable for food-safety monitoring.

