## Supplementary tables for "FRET-based aptamer assay for sensitive detection of *Salmonella paratyphi* A and revealing its molecular interaction with DNA gyrase"

| **S.NO** | **Samples** | **Mean Pore diameter (nm)** | **Properties BET surface area/m^2^/g** | **Pore volume / cm^3^/g** |
| --- | --- | --- | --- | --- |
| 1 | Graphene | 6.1 | 266.9 | 1.48 |
| 2 | Graphene oxide | 8.835 \| \| | 221.2 | 0.97 |

| **Aptamer** | **Position** | **Length** | **QGRS** | **G-Score** |
| --- | --- | --- | --- | --- |
| **Sal1** | 15 | 19 | GGGCCGCGGGTCAGGGGGG | 68 |
| **Sal2** | 19 | 15 | GGTGGGTTTGGCCGG | 35 |

**Supplementary Table 1: Predicted G score using QGRS mapper**

**Supplementary Table 2: BET surface analysis of synthesised GO**
