## Supplementary Figures for "FRET-based aptamer assay for sensitive detection of *Salmonella paratyphi* A and revealing its molecular interaction with DNA gyrase"

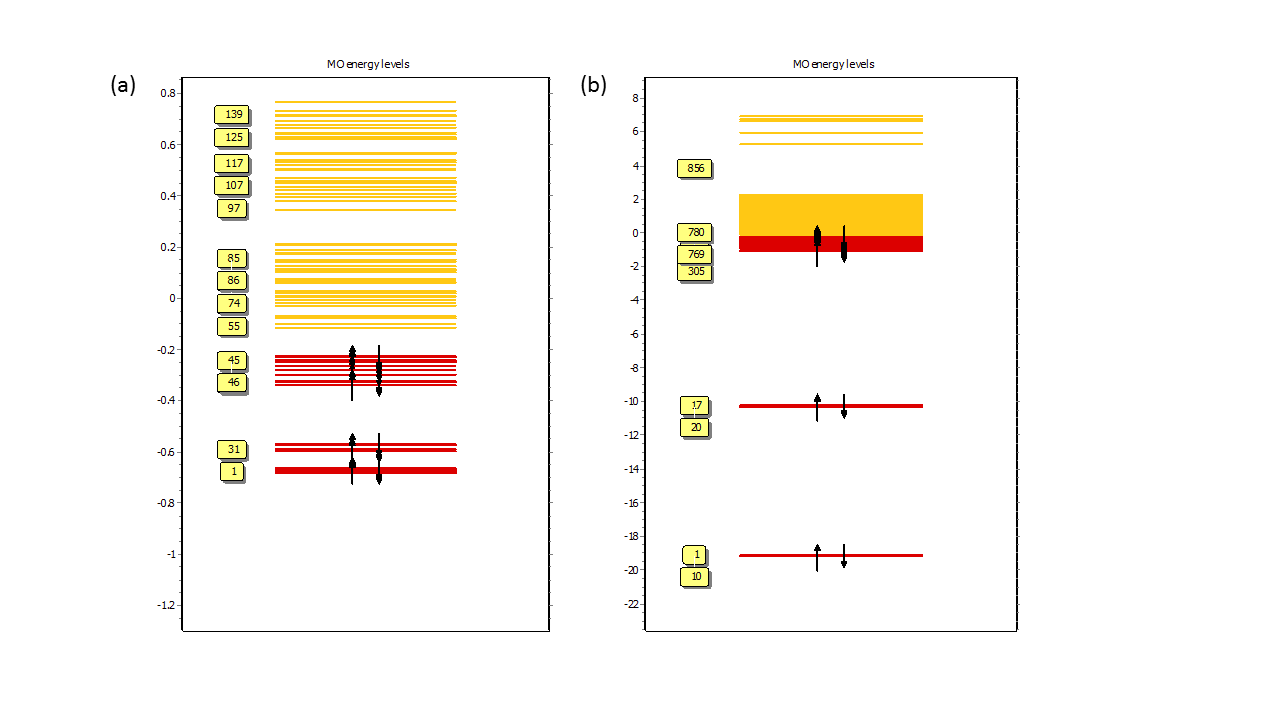

**Supplementary Figure 1**: Molecular orbital energy levels of (a) Quantum dot and (b) graphene oxide conjugated	quantum dot.

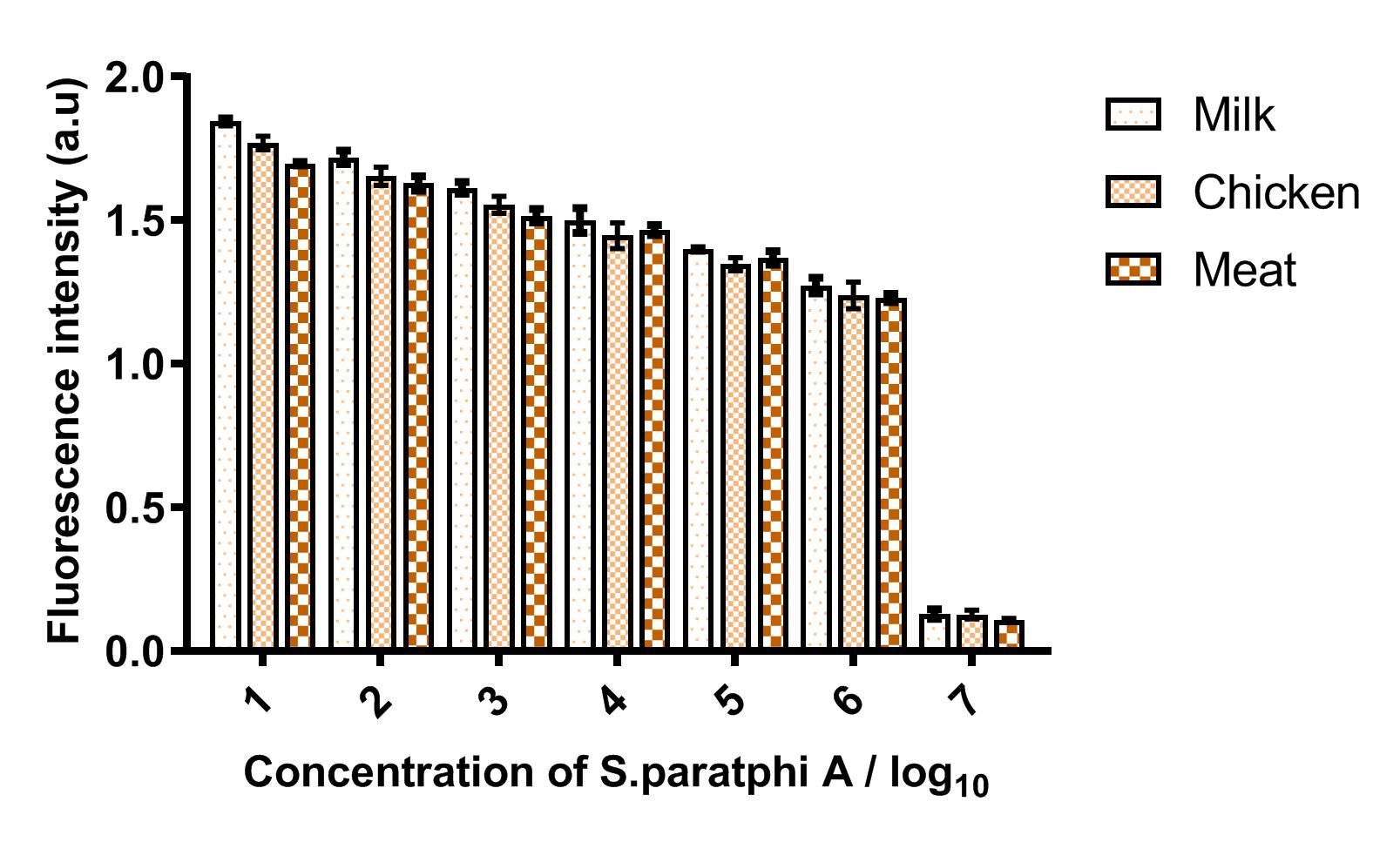

**Supplementary Figure 2: Matrix interference of developed FRET based aptasensor**

Matrix interference assay for various samples spiked with *Salmonella paratyphi A.* Comparison study yieled a P value of <0.001 for independent experiments performed and statistically evaluated using ANOVA.

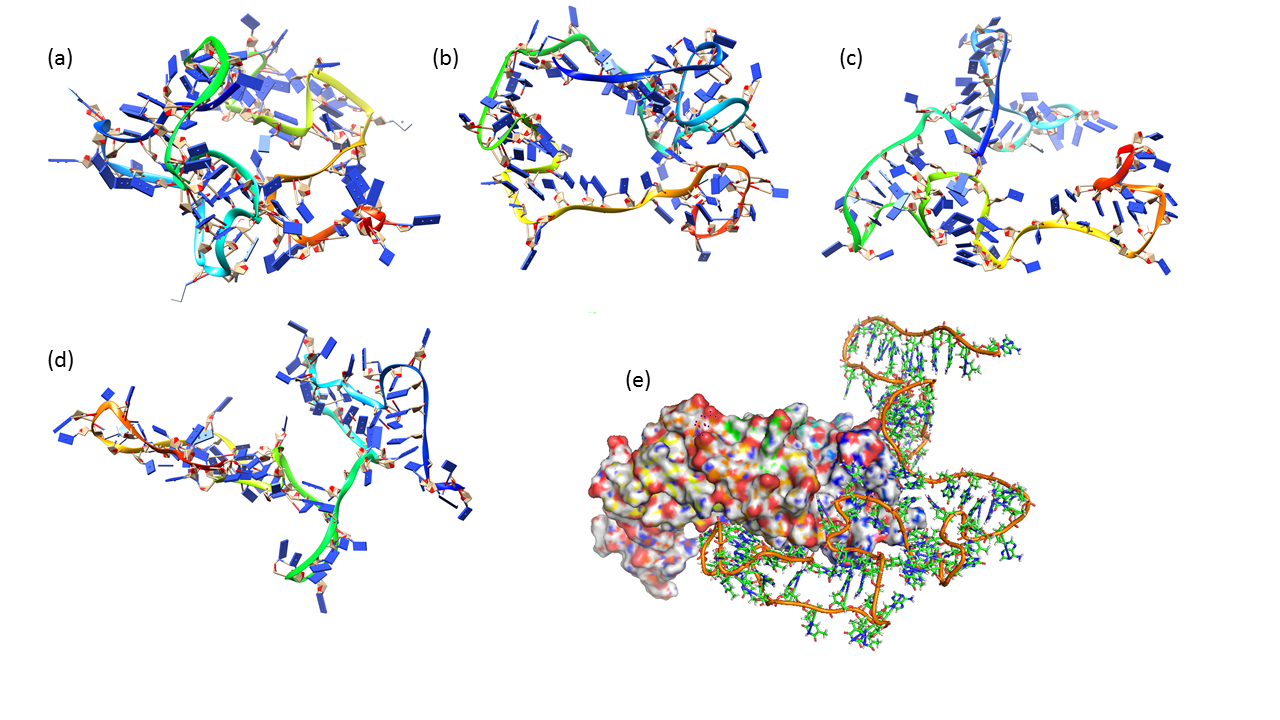

**Supplementary Figure 3:** (a) - (d) time evolution of ssDNA three dimensional structure (e) represents the binding pose of ssDNA 3-D structure of DNA-gyrase protein.

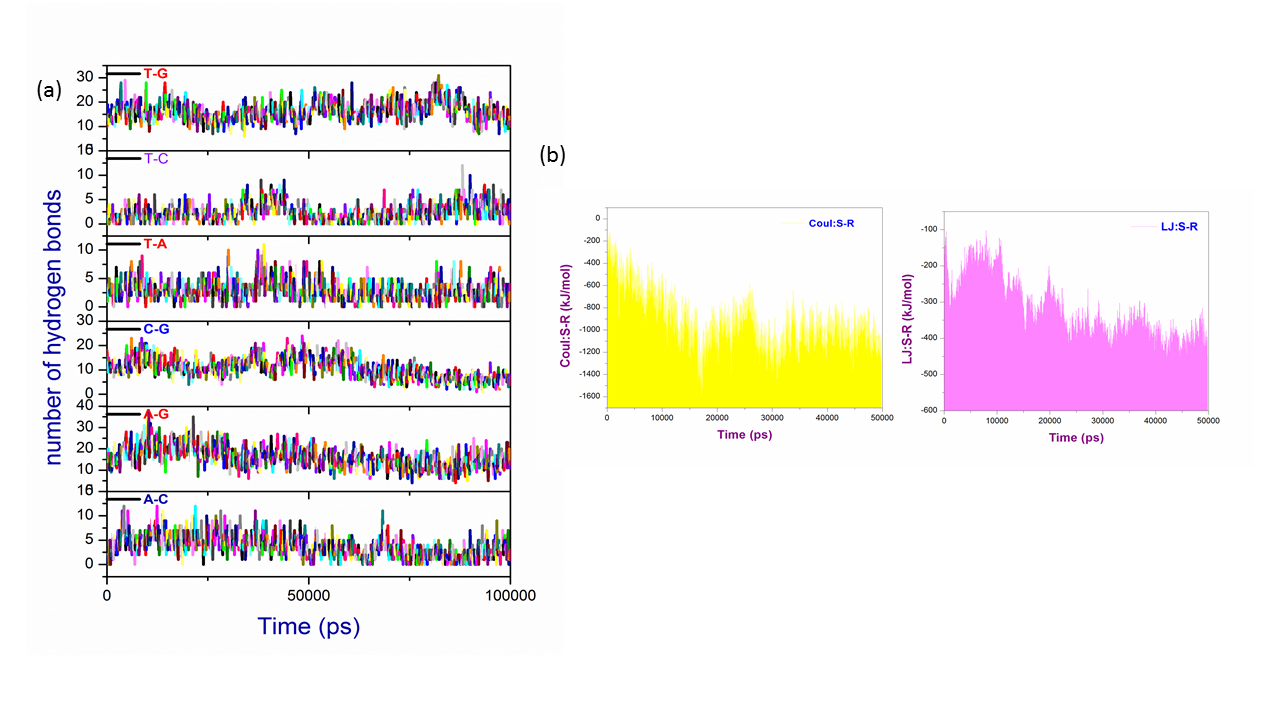

**Supplementary Figure 4**: (a) The number of hydrogen bond formed between individual pairs of the ssDNA and (b) non-bonded interaction energy between ssDNA and DNA-gyrase protein

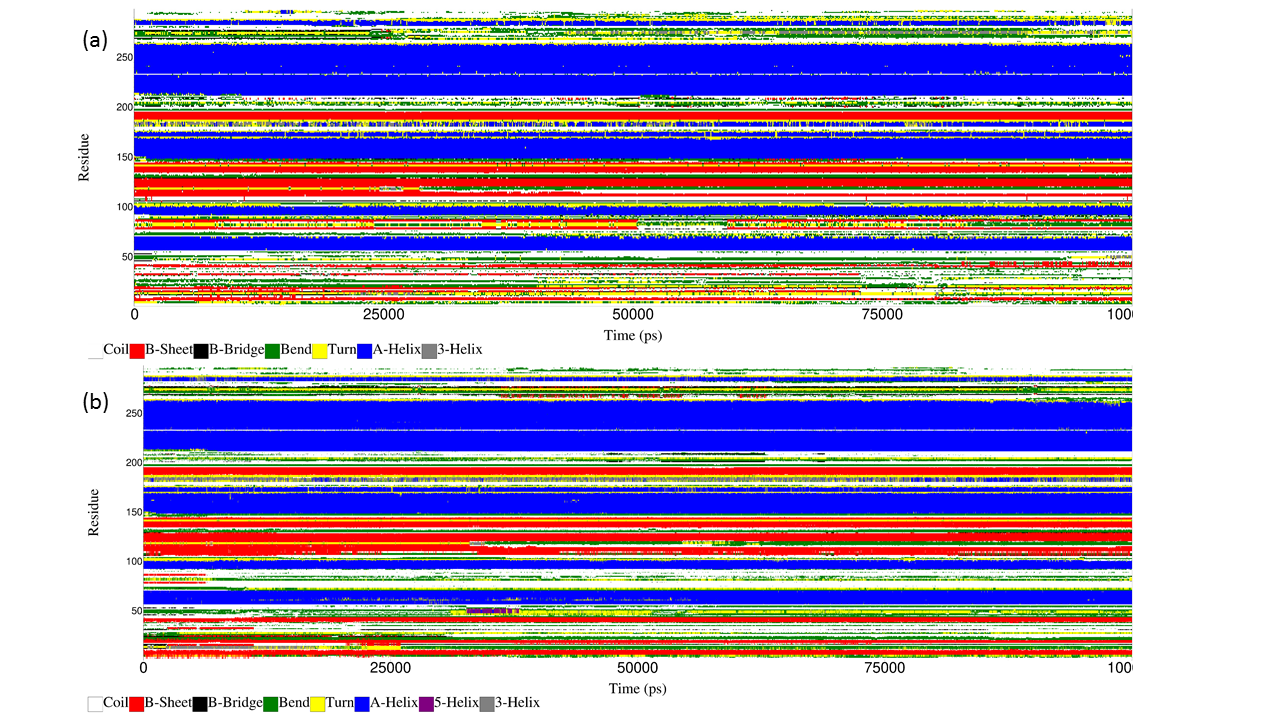

**Supplementary Figure 5:** The time evolution of secondary structure DNA-gyrase protein (a) native protein (b) with ssDNA.
